# A very long chain fatty acid responsive transcription factor, MYB93, regulates lateral root development in Arabidopsis

**DOI:** 10.1101/2022.05.24.493307

**Authors:** Yuta Uemura, Saori Kimura, Tomomichi Ohta, Takamasa Suzuki, Kosuke Mase, Hiroyuki Kato, Satomi Sakaoka, Yuki Komine, Kazuhiro Hotta, Motoyuki Shimizu, Atsushi Morikami, Hironaka Tsukagoshi

## Abstract

Lateral roots (LRs) are critical to rhizosphere development in plants. Although the molecular mechanisms by which auxin regulates LR development has been studied extensively, many additional regulatory systems are thought to be involved. Based on the expression analysis of LTPG1 and 2, we found that they were specifically expressed at the developing LR primordium (LRP), and the number of LRs were reduced in these mutants. Because LTPG is a protein that transports Very Long Chain Fatty Acids (VLCFAs), we hypothesized that VLCFAs regulated LR development. We revealed that late LRP development was stunted when VLCFA levels were reduced in the synthetic mutant, *kcs1-5*. In this study, we established a method to analyze the LRP development stages with high temporal resolution using a deep neural network. We identified a VLCFA responsible transcription factor, MYB93, from transcriptome analysis of *kcs1-5*. Interestingly, MYB93 showed a carbon chain length-specific expression response after treatment of VLCFA. Furthermore, *myb93* transcriptome analysis suggested that MYB93 regulated the expression of cell wall organization genes. Our results indicated that VLCFA is a regulator of LRP development through transcription factor-mediated regulation of gene expression.

## Introduction

Robust development of the rhizosphere, which comprises primary root and lateral root (LR) systems, is an important process in the growth of plants due to their sessile nature. The location and timing of LR primordium (LRP) development and LR emergence is controlled by a very precise auxin-related mechanism that has been extensively analyzed at the molecular level (Parizot, et al, 2007; Dubrovsky et al., 2008; Du and Scheres, 2018; Motte and Beeckman, 2019). At the later stages of LR development, the LRP crosses the lateral root overlying cells (LROCs; endodermis, cortex, and epidermis cells) of the primary root. Communication between LROCs and LRPs is crucial for the emergence of LRs (Teixeria and ten Tusscher, 2019). In this event, auxin transport in LROCs is essential for allowing LRP passage (Swarup et al., 2008; Kumpf et al., 2013; Vermeer et al., 2014). However, studies have suggested that other factors of auxin are involved in LR development. For example, it has been reported that very long chain fatty acids (VLCFAs) are involved in LR development and pericycle competence for root callus formation (Shang et al., 2016; Trinh et al., 2019; Lv et al., 2021).

VLCFAs are important molecules in the plant life cycle. VLCFAs are critical components of biological membranes, and changes to lipid composition can affect membrane properties. (Devaiah et al., 2006; Markham et al., 2013; Batsale, et al., 2021). VLCFAs are considered 20 or more carbon chains length fatty acids and are synthesized in the endoplasmic reticulum in plants (Bach and Faure., 2010). Four enzyme complexes, ß-ketoacyl-CoA synthase (KCS), ß-ketoacyl-CoA reductase (KCR), ß-hydroxyacyl-CoA dehydratase (HCD), and enoyl-CoA reductase (ECR), are involved in the biosynthesis of VLCFAs (Millar and Kunst, 1997; Zheng et al., 2005; Bach et al., 2008; Joubès et al., 2008; Beaudoin et al., 2009; Morineau et al., 2016). Several loss-of-function VLCFA mutants show embryo lethality and strong growth defects (Bach et al., 2008; Beaudoin et al., 2009). In addition, VLCFAs act as a signal molecule that regulates development of shoot apical meristem with non-cell autonomous manner (Nobusawa et al., 2013). Moreover, VLCFAs are involved in cell cytokinesis, endocytosis, and signal transduction responses to abiotic, biotic-stresses (Roudier et al., 2010; Bach et al., 2011; Molino et al., 2014; De Bigault Du Granrut and Cacas, 2016). The transport of VLCFAs to the cell surface is also an important regulatory system in plant development. Among putative transporters, GPI-anchored lipid transfer proteins (LTPGs) are well characterized. *LTPG1* and *LTPG2* are involved in stem cuticle wax transport (DeBono et al., 2009; Kim et al., 2012). Other LTPG genes are involved in seed coat suberization and pollen development (Edstam and Edqvist, 2014). Moreover, *LTPG1* and *LTPG2* are involved in root cell elongation in response to hydrogen peroxide (Mabuchi et al., 2018).

Recent studies reveal transcriptional regulators of VLCFA biosynthesis genes. An AP2/EREBP family transcription factor, PUCHI, regulates the expression of various VLCFA synthesis genes, and *puchi-1* mutants exhibit reduced VLCFA levels and delayed LR development (Trinh et al., 2019). Another AP2/EREBP family transcription factor, ERF13, regulates LRP emergence by controlling the expression of *KCS16* (Lv et al., 2021). In addition to transcriptional regulation, a cuticularized outermost layer called the root cap cuticle (RCC), containing VLCFAs, forms around the LRP and contributes to the emergence of normal LRs (Berhin et al., 2019). However, the molecular mechanisms of VLCFA regulating transcriptional network that controls LR development have not been elucidated.

In this study, we found that *LTPG1* and *LTPG2* were specifically expressed in the developing LRP, prompting us to investigate the role of VLCFAs in LR development. Analysis with deep neural network (DNN) machine learning, which was able to follow continuous LR development, indicated that VLCFA regulated later stages of LR development. Moreover, we identified a VLCFA transcription factor, MYB93, from transcriptome analysis of the *kcs1-5* VLCFA synthesis mutant. We found that MYB93 controlled late-stage LR development. Transcriptome analysis showed that MYB93 regulated a set of cell wall remodeling related genes, including expansins. Expression analysis indicated that MYB93 regulated its downstream genes in LROCs. These results suggested that VLCFAs regulated LR development by modulating cell wall remodeling through the expression of *MYB93* in LROCs. Additionally, we found that extracellular transport of VLCFAs by LTPG1 and LTPG2 was involved in LR development by contributing to RCC formation. Our results identified VLCFA as an important compound involved in various stages of LR development.

## Results

### LTPG1 and LTPG2 are involved in LR development

We investigated *LTPG1* and *LTPG2* expression patterns in roots using the translational fusions of LTPG1 and LTPG2 (Figure 1A). Notably, both *LTPG1* and *LTPG2* were weakly expressed in the LRP, starting from stage I. LTPG1 was expressed in the LRP during stages I, II, and III, and its expression became weaker through stage IV. LTPG2 expression continued until the later stage of LRP development. Interestingly, expression of both genes was limited in the outermost cell layer of the LRP. This indicated that LTPG1 and LTPG2 were involved in LR development.

**Figure 1.**
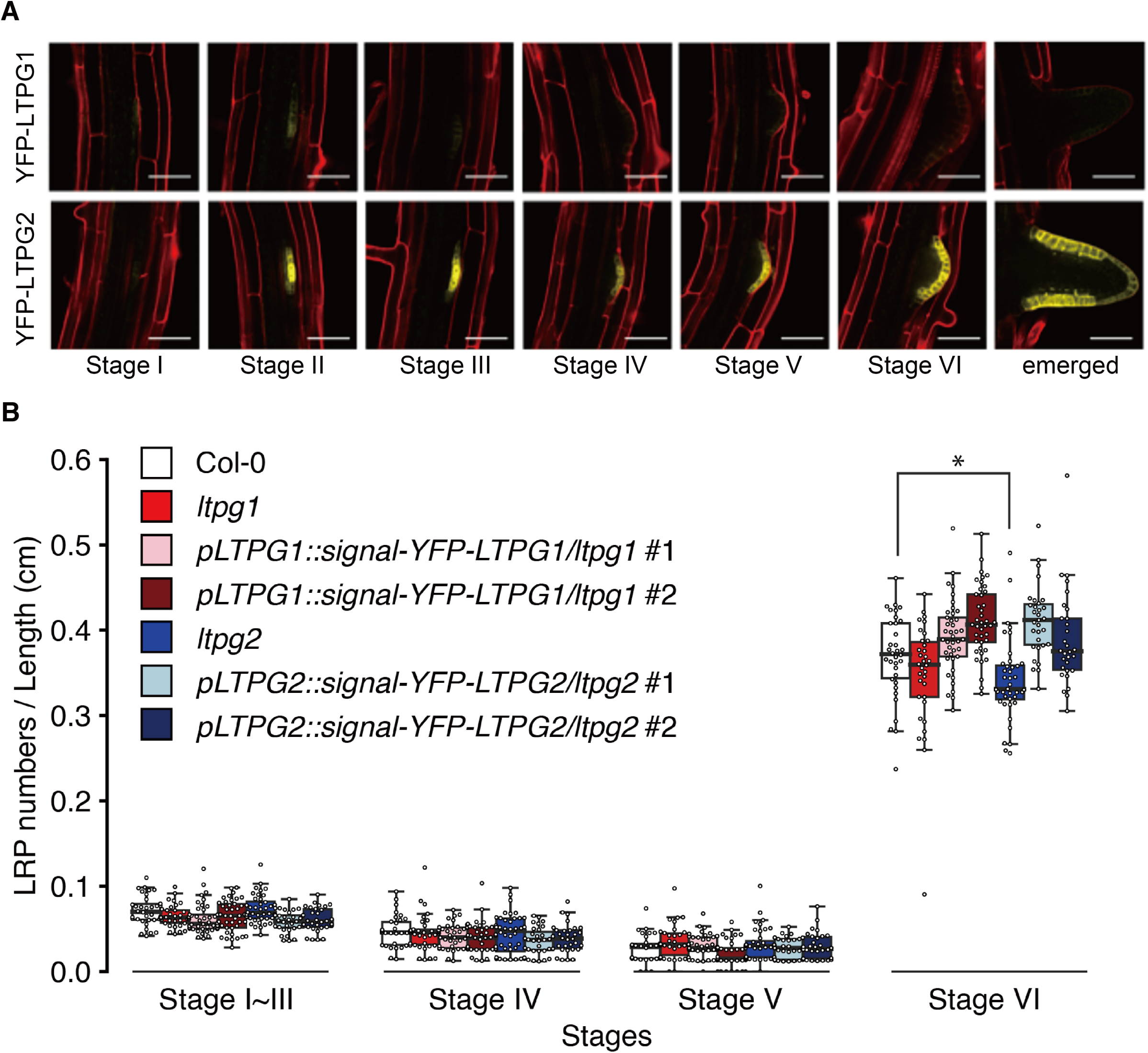
LTPG1 and LTPG2 are expressed in the lateral root primordium and regulate lateral root development. A, *pLTPG1::YFP-LTPG1* and *pLTPG2::YFP-LTPG2* expression in different stages of lateral root development. Roots were stained with propidium iodide (PI). Scale bars, 50 µm. B, Distribution of lateral root development in 13-day-old seedlings of Col-0, *ltpg1*, and *ltpg2* plants, and *ltpg1* and *ltpg2* complemented lines with *pLTPG1::YFP-LTPG1* and *pLTPG2::YFP-LTPG2* (n = 30). Significant differences from Col-0 of each lateral root stage were detected by Student’s *t*-test (\**p* < 0.05).

To test this possibility, we counted LR numbers in *ltpg1* and *ltpg2* mutants and compared to wild type (Col-0) roots (Figure 1B). *ltpg2* mutants exhibited decreased numbers of stage VI emerged LRs. *ltpg1* also decreased number of stage VI, however there was no significant change in the number of LRPs in *ltpg1* compared to the wild type. This might be due to the lower expression level of *LTPG1* at LRP compared to *LTPG2* (Figure 1A). The LR phenotypes of *ltpg1* and *ltpg2* were complimented with translational fusions of LTPG1 and LTPG2 in each mutant (Figure 1B). These results indicated that LTPG2 was involved in LR development.

### VLCFAs regulated LR development

To test VLCFAs effects on LR development, we treated roots with the VLCFAs synthesis inhibitor, cafenstrole (300 µM). This chemical is known as inhibitor of KCS activity, which adds C2 from malonyl-CoA to the fatty acid substrates (Trenkamp et al., 2004; Shang et al., 2016). Cafenstrole treatment increased stage V and reduced stage VI LRP counts significantly (Figure 2A). This result indicated that reduced VLCFA levels inhibited later stage LRP development.

**Figure 2.**
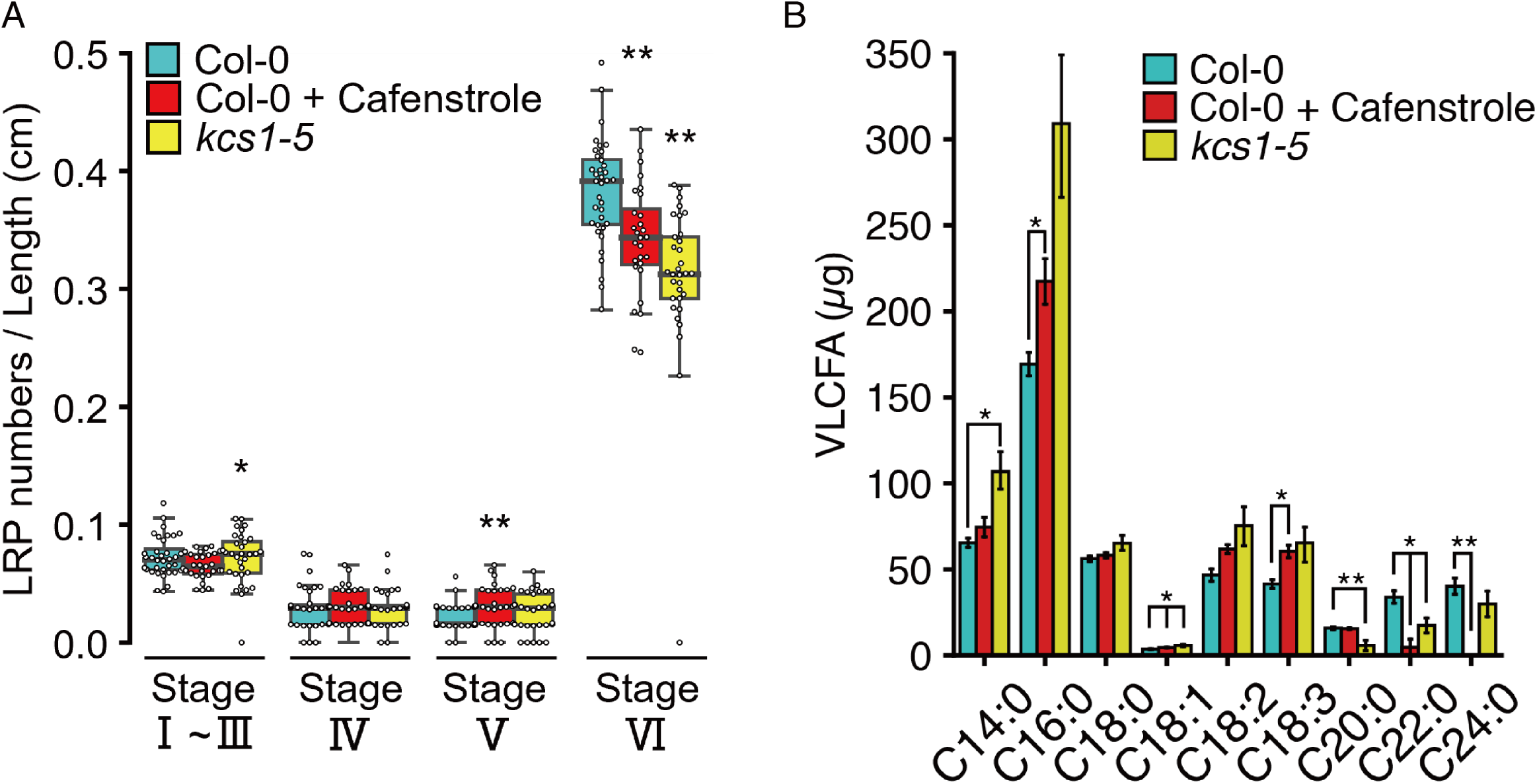
Alteration of VLCFA levels affects lateral root development. A, Distribution of lateral root development stages in 13-day-old seedlings of Col-0, Col-0 treated with cafenstrole (300 nM), and *kcs1-5* (n = 30). Significant differences from Col-0 were detected by Student’s *t*-test (\*\**p* < 0.01, \**p* < 0.05). B, Fatty acids composition of 13-day-old seedlings of Col-0, Col-0 treated with cafenstrole (300 nM), and *kcs1-5*. Data are represented as means ± SD of three biological replicates. Significant differences from Col-0 of each stage were detected by Student’s *t*-test (\*\**p* < 0.01, \**p* < 0.05).

We also tested the effects of VLCFAs on LR development genetically using *kcs1-5* VLCFA synthesis enzyme mutants. KCS1 catalyzes the reaction for elongation of VLCFA from long chain fatty acid (Millar et al., 1997; Todd, et al., 1999). In *kcs1-5* mutants, the numbers of stage I - III LRs significantly increased, and the number of stage VI was significantly reduced as seen under cafenstrole treatment (Figure 2A). These results supported the regulatory role of VLCFA in later-stage LR development.

We measured fatty acids contents in Col-0 under control and cafenstrole treatment, and in *kcs1-5* mutants (Figure 2B). In *ksc1-5*, C20, C22, and C24 fatty acid contents significantly decreased comparing with Col-0 under control condition. Col-0 treated with cafenstrole also exhibited significantly reduced levels of C22 and C24. These results strongly indicated that KCS1 catalyzed fatty acid elongation from C18 to C20 in roots, whereas cafensterole inhibited the subsequent reaction from C20 to C22. We also measured fatty acids contents in *ltpg1* and *ltpg2* mutants and did not observe any significant differences of VLCFA contents between *ltpg1*, or *ltpg2* mutants and Col-0 (Supplemental Figure S1). Based on the phenotype of LR abundance in *kcs1-5*, *ltpg1,* and *ltpg2* mutants, both VLCFA levels and transport by LTPGs were important for LR development.

### VLCFA affected late-stage LR development

To more closely investigate the role of VLCFAs in LR development, we induced LRP formation using a gravitropic stimulus assay (Figure 3A). After 20 h gravistimulation, Col-0 with cafenstrole treatment and *kcs1-5* exhibited an increased number of stage I LRPs and reduced number of stage II LRPs. After 32 h, the number stage I and II LRPs in Col-0 under cafenstrole treatment and in *kcs1-5* was higher than in Col-0 under control condition, and the number of stage III and IV LRPs was considerably lower in cafenstrole-treated Col-0 plants than in Col-0 under control condition. In *kcs1-5*, no change was seen in the abundance of stage III LRPs, but the number of stage IV LRPs decreased significantly. After 46 h, the number of LRPs in stages II and III remained high in roots of cafenstrole-treated Col-0 and *kcs1-5*. However, in the cafenstrole-treated Col-0 roots, no change was seen in the number of stage V LRPs, but the number of LRP stage VI LRPs decreased dramatically. In *kcs1-5,* the number of stage IV LRPs increased, and the number of stage V and stage VI LRPs decreased, significantly so for stage VI. These effects were partially ameliorated by exogenous VLCFA mix (C20, C22, and C24) treatment in both cafenstrole- treated Col-0 and *kcs1-5* roots, especially for stage IV LRPs at 32 h and stage VI at 46 h. At 20 h post-gravistimulation, the effect of exogenous VLCFA was hardly observed. These results strongly indicated that VLCFA played an important role in later LRP development and emergences, rather than in LRP formation.

**Figure 3.**
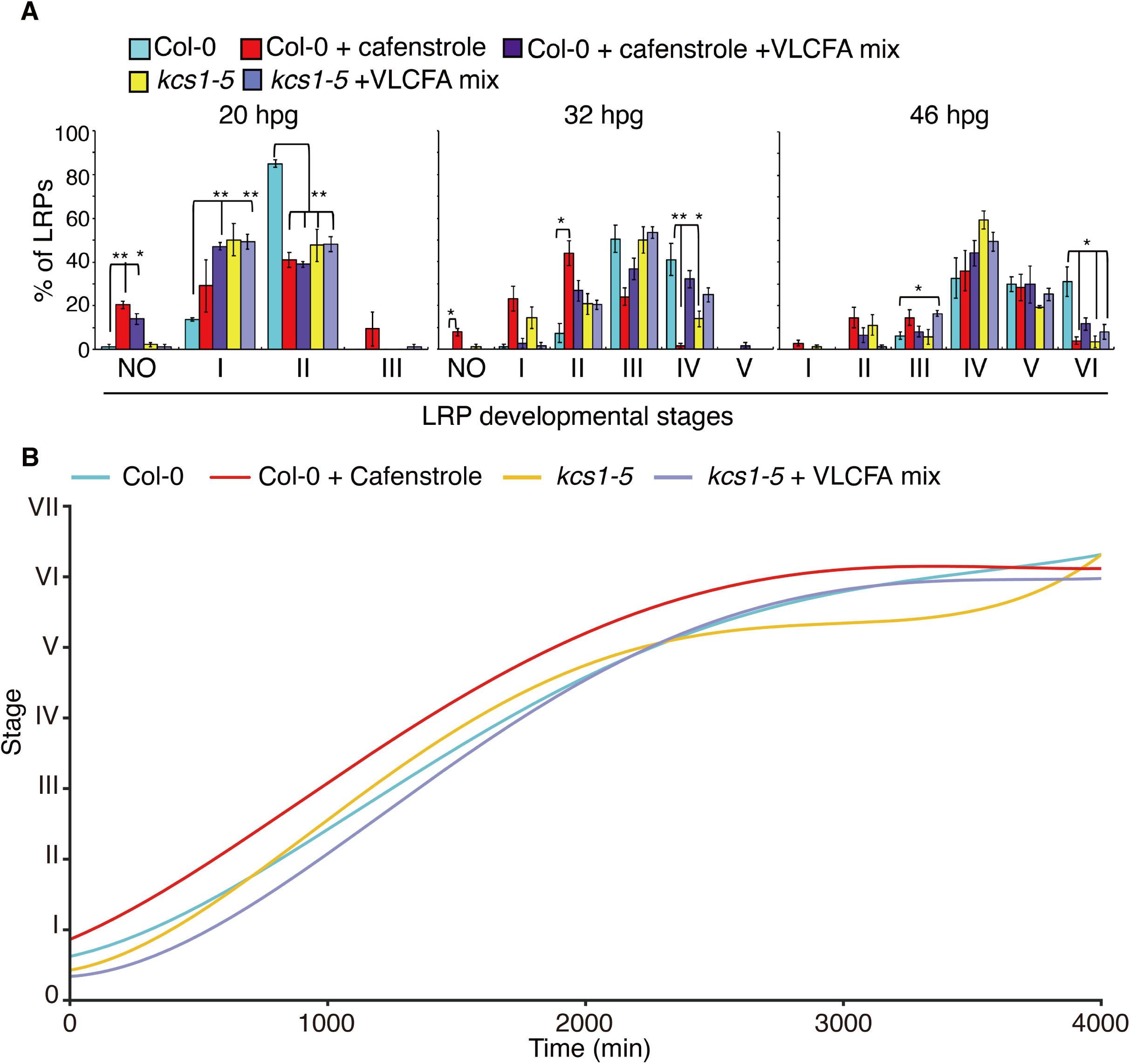
VLCFA is involved in later stages of lateral root development. A, Stages of lateral root primordia development after 20 h, 32 h, and 46 h of gravistimulation were classified in Col-0; Col-0 treated with cafenstrole (300 nM); Col-0 treated with cafenstrole (300 nM) and VLCFA mix (200 µM each of C20, C22, and C24); *kcs1-5*; and *kcs1-5* treated with VLCFA mix (200 µM each). Stages were determined as described by Malamy and Benfey (1997). Data are represented as mean ± SE of three biological replicates, with 20 seedlings in each. Significant differences in number of lateral roots of each stage from Col-0 were detected by Student’s *t*-test (\*\**p* < 0.01, \**p* < 0.05). B, Measurement of lateral root stage transition time for Col-0 (light blue); Col-0 treated with cafenstrole (300 nM; red); *kcs1-5* (yellow); and *kcs1-5* treated with VLCFA mix (200 µM each; purple) by DNN (n = 29, 19, 24, and 12, respectively). Generated timeseries data of lateral root primordium stages were approximated as a quadratic function.

Next, we conducted time-lapse imaging of LRP development and analyzed LRP developmental patterns using Deep Neural Network (DNN) imaging analysis (Figure 3B). We used 3,923 images of LRPs to train DNN learning, then the accuracy of DNN stage classification compared with manual classification. The difference in mean absolute errors (MAEs) between human and DNN classification was not significant (*p* = 0.3887; Supplemental Figure S2C). With our classification of LRP stages in DNN, we can recognize more images at a time than a human classification. In other words, it is possible to perform LRP stage classification of time lapse images, which improves the temporal resolution of LRP development analysis. This analysis also can be tracking the development of a single LRP and estimate the pattern of LRP development. Next, we captured time lapse images from Col-0, Col-0 treated with cafenstrole, *kcs1-5*, and *kcs1-5* treated with VLCFAs every 30 min for 72 h and analyzed them with our DNN. Analysis showed that length of transition time from LRP stage I to V was not different between Col-0 and *kcs1-5*. However, the transition from LRP stage V to VI was longer in *kcs1-5* than in Col-0, and the approximate growth curve of *kcs1-5* was stagnant between stage V and VI. Similarly, the developmental speed of the wild type LRP in cafenstrole-treated Col-0 roots was also stagnant at late LRP stages. This effect was attenuated in *kcs1-5* by VLCFA mix treatment, which resulted in control-level growth. Our DNN analysis strongly supported the hypothesis that VLCFAs regulate the transition of late-stage LRP development.

### Reduced VLCFA levels affected the expression of several genes

To elucidate how a reduction in VLCFA levels affected gene expression regarding LR development, we conducted RNAseq analysis and compared whole-root gene expression between Col-0 and *kcs1-5* mutants. A total of 60 and 8 genes (singletons) were significantly upregulated and downregulated, respectively, between the two groups (two fold change, FDR < 0.05; Supplemental Data Set S1). We analyzed the Gene Ontology (GO) terms of “Biological Process” and “Molecular Functions” of upregulated genes in *ksc1-5* mutants, and found that 13 “Biological Process” terms were significantly enriched (*p* value < 0.001; Figure 4, A and B). Interestingly, auxin related GO terms were not enriched in our data set. Cell wall-related GO terms, such as “phenylpropanoid metabolic process,” “polysaccharide metabolic process,” and “cell wall organization or biogenesis” were enriched, indicating that lowered VLCFA levels in *kcs1-5* affected cell wall status. Notably, “regulation of transcription from RNA polymerase II promoter” and “transcription factor activities” terms were strongly enriched, indicating gene regulatory network responses to altered VLCFA levels (Figure 4B). In fact, a 5 MYB domain containing transcription factor genes was upregulated in *kcs1-5*. These results suggested that the existence of a gene regulatory network controlled by transcription factors in response to VLCFAs.

**Figure 4.**
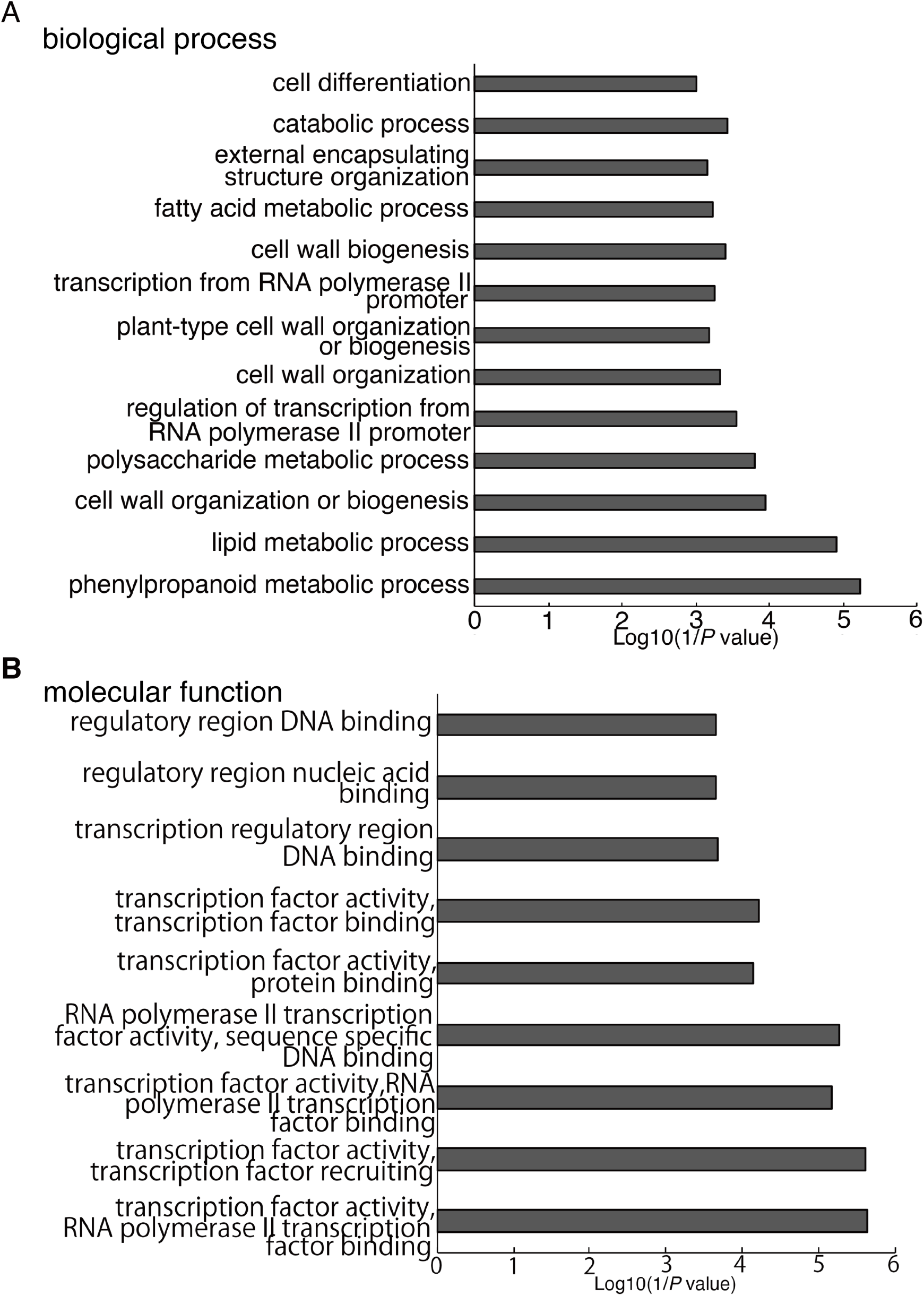
Differentially expressed genes in *kcs1-5* mutant roots. (A and B) Gene Ontology terms related to “Biological Process” (A) and “Molecular Function” (B) enriched among genes significantly upregulated genes in *kcs1-5* roots comparing to Col-0.

### The transcription factor MYB93 responded to VLCFAs to regulate LR development

Among the differentially expressed genes in *ksc1-5*, we identified one encoding the transcription factor, MYB93. We used quantitative real-time RT-PCR (RT-qPCR) to examine *MYB93* expression in response to VLCFA. *MYB93* expression level was significantly higher in *kcs1-5* roots than in Col-0 (Figure 5A). Moreover, *MYB93* expression in *kcs1-5* was reduced to control levels after treatment with VLCFA mix, C20, C22, and C24 treatment for 24 h in *kcs1-5*. Interestingly, C18 treatment for 24 h didn’t affect *MYB93* expression (Figure 5A). Then, we performed a time course RT-qPCR analysis in Col-0 and *kcs1-5* treated with VLCFA mix. MYB93 was suppressed to the same level as Col-0 in *kcs1-5* treated with VLCFA mix for 3 h and 6 h (Figure 5B). These results indicated that *MYB93* was a VLCFA-responsive transcription factor. Next, we investigated *MYB93* expression patterns using cyan fluorescent protein (CFP) translational fusion of MYB93 (*pMYB93::CFP-MYB93*). CFP-MYB93 was detected specifically in the endodermis of LROCs (Figure 5C). *MYB93* expression was first observed in stage I of LROCs, and increased until stage V (Figure 5C). These results indicated that MYB93 was expressed specifically LROCs, but not in LRPs.

**Figure 5.**
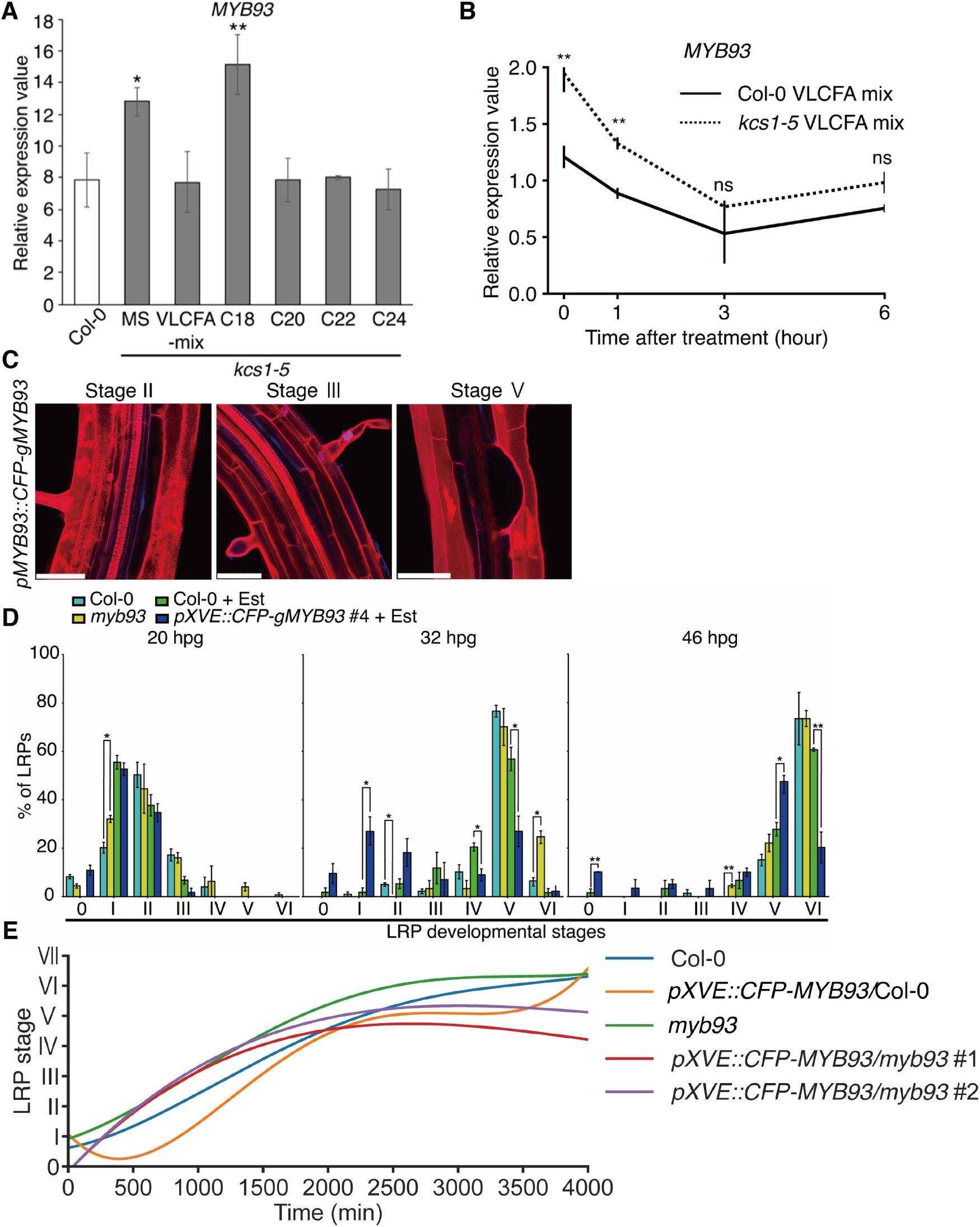
The VLCFA responsive transcription factor MYB93 regulates late-stage lateral root development. A, RT-qPCR analysis of *MYB93* in whole roots of Col-0; *kcs1-5*; and *kcs1-5* treated with VLCFA mix (200 µM each C20, C22, and C24); C18 (200 µM); C20 (200 µM); C22 (200 µM); or C24 (200 µM) (n = 3; ±SD). Significant differences from Col-0 were detected by Student’s *t*-test (\*\**p* < 0.01, \**p* < 0.05). B, Time course RT-qPCR analysis for the expression of *MYB93* after 0, 1, 3, or 6 h of VLCFA mix treatment in the root of Col-0 (solid line) and *ksc1-5* (dashed line). (n = 3; ±SD). Significant differences from Col-0 were detected by Student’s *t*-test (\*\**p* < 0.01). C, Expression of *pMYB93::CFP-gMYB93* in the lateral root primordia of Col-0 background. Roots were stained with PI. Scale bars, 50 µm. D, Stages of lateral root primordia after 20, 32, and 46 h of gravistimulation in Col-0, *myb93*, Col-0 treated with 5 µM estradiol, and *pXVE::CFP-MYB93* roots treated with 5 µM estradiol. Data are represented as mean ± SE of three biological replicates, with 20 seedlings in each. Significant differences from Col-0 or Col-0 treated with estradiol were detected by Student’s *t*-test (\*\**p* < 0.01, \**p* < 0.05). E, DNN measurement of lateral root stage transition time for Col-0 (blue), *pMYB93::CFP-gMYB93*/Col-0 (orange), *myb93* (green), *pMYB93::CFP-gMYB93*/*myb93* #1 (red), and *pMYB93::CFP-gMYB93*/*myb93* #2 (purple) plants treated with 5 µM estradiol (n = 29, 10, 14, 12, and 14, respectively: Col-0 was the same data as Figure 3B). Generated time series data of lateral root primordium stages ware approximated as a quadratic function.

We used gravistimulation to identify which stage of LR development was regulated by MYB93 (Figure 5D). After 20 h gravistimulation, in *myb93* the number of stage I, V, and VI LRPs increased significantly. After 32 h, the number of LRPs in stage VI had increased significantly in *myb93*. After 46 h, there was no change in the number of stage V or VI LRPs, but the number of LRPs over stage VI increased significantly. To investigate the effects of *MYB93* overexpression, we constructed an estradiol-inducible version of MYB93 fused with CFP (Supplemental Figure S3). We then treated the estradiol-inducible CFP-MYB93 plants (*pXVE::CFP-MYB93*) with gravistimulation after estradiol treatment for 24 h. In this assay, we used estradiol-treated Col-0 as the control for comparison. Twenty hours after gravistimulation, *pXVE::CFP-MYB93* plants exhibited a decreased number of LRP stage III roots, but the difference was not significant. After 32 h, number of stage 0, I and II LRPs had increased (significantly so for stage I), whereas the number of stage IV and V LRPs decreased significantly. This result indicated that MYB93 induction delayed LRP development. After 46 h, the number of stage 0, I, II, III, and V LRPs had increased (significantly so for stages 0 and V), and that of stage VI LRPs had decreased significantly. These results indicated that MYB93 had an inhibitory effect on the later stages of LRP development. We also found that estradiol itself affected the LRP development in Col-0 plants, though not in the same manner as MYB93. In particular, the number of stage I LRPs increased at 20 h post-gravistimulation, and that of stage IV LRPs increased at 32 h, while numbers of stages V and VI LRPs decreased at 32 h. Furthermore, at 46 h, a significant increase in stage V LRPs was observed. Although estradiol affected LRP development in Col-0 plants, the late stage of development was more considerably delayed in *pXVE::CFP-MYB93* roots.

In addition to manually LRP counting, we performed a DNN analysis using time-lapse images to quantify the transition of LRP development (Figure 5E). In *myb93* mutants, LR development proceeded faster than in the wild type plants starting around stage II, and the time needed to reach stage VI LR emergence was about 800 minutes shorter. On the contrary, in estradiol-induced *pXVE::CFP-MYB93* plants, LR development progressed more slowly from stage II than in wild type plants, and stages V and VI exhibited much slower development and transition than in wild type plants. This developmental curve was very similar to that of *kcs1-5* mutants, and was consistent with the upregulated expression of *MYB93* in *kcs1-5*. In addition, overexpression of *MYB93* in *myb93* mutants (Supplemental Figure S3) showed delayed LR development from stage IV in two independent lines (Figure 5E). This delayed LR development was the opposite phenotype of *myb93* mutants, indicating that MYB93 regulated the development of late LR stages.

### MYB93 regulates cell wall organization genes and controls LR development

To explore the expression pattern of *MYB93* regulated genes during LRP development, we conducted RNAseq analysis using only the LRP developing sites of roots gravistimulated for 46 h to explore the set of genes regulated by MYB93. As a result of RNAseq, 88 and 10 genes (singletons) were significantly upregulated and downregulated (excluding *MYB93* itself), respectively, in *myb93* mutant (two-fold change, FDR < 0.05; Supplemental Data Set S2). We analyzed GO terms under “Biological Process” for the upregulated genes, and found that 50 GO terms were significantly enriched (*p* value < 0.001). Among them, terms related to cell wall biogenesis and organization showed remarkable enrichment (Figure 6A). This group included genes for regulatory enzymes of cell wall remodeling and cell-cell adhesion, such as extensin-like proteins, expansins, peroxidases, and pectin lyase families. Among the expansins, *EXPA17*, which has been reported to be involved in LR development (Lee and Kim, 2013), was significantly upregulated in the *myb93* RNAseq data set. In addition, *EXPA7* and *EXPA18* were upregulated in *myb93* mutants. We confirmed the expression of these three expansin genes at the site of gravistimulation by RT-qPCR. Only EXPA17 showed strong upregulation in the *myb93* (Figure 6B). Next, we constructed promoter GFP reporter lines of the MYB93 downstream expansin genes. In the root tips, these expansins were expressed in epidermis of the elongation and the maturation zones of roots (Supplemental Figure S4). In addition, *pEXPA17::GFP* plants showed specific expression in LROC cortex cells. *pEXPA7::GFP* and *pEXPA18::GFP* were also expressed in LROCs, in epidermis cells instead of in cortex (Figure 6C). These expression patterns indicated that the expansins regulated LR development from LROCs. We further investigated the expression levels of these putative MYB93-regulated expansin genes in *pXVE::CFP-MYB93* plants, and found that expression was suppressed by estradiol treatment. We also found that expression of these genes was suppressed in two independent *pXVE::CFP-MYB93* in *myb93* mutant background significantly (Figure 6D). These results suggested that MYB93 was a transcriptional repressor of expansins and controlled late-stage LRP development by repressing expansin gene expression, especially that of *EXPA17*, in LROCs.

**Figure 6.**
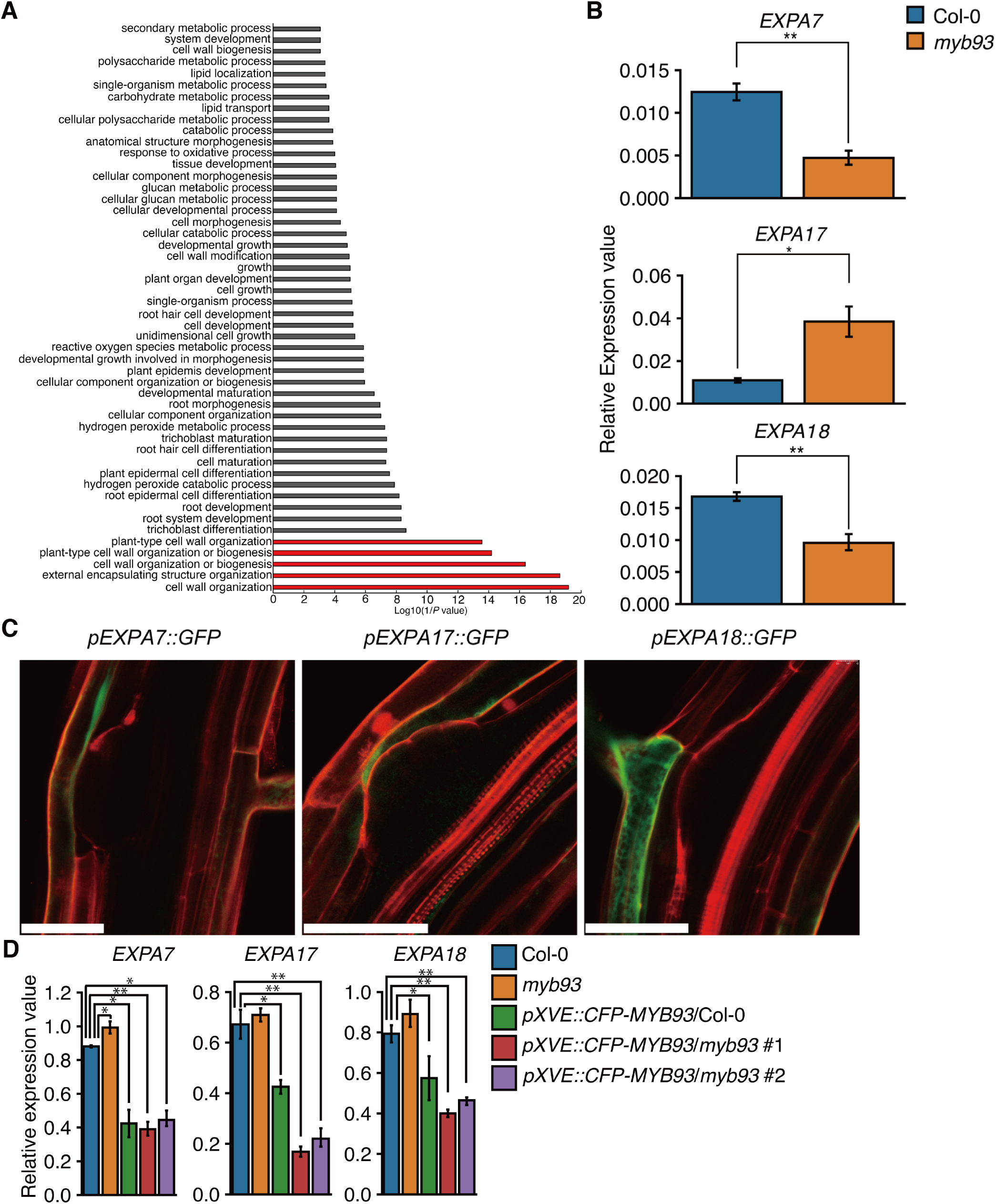
MYB93-regulated genes. A, Gene Ontology terms related to “Biological Process” enriched among significantly upregulated genes in *myb93* mutant roots. B, RT-qPCR analysis of MYB93 regulated genes after 46 h of bending site gravistimulation in Col-0 and *myb93* plants (n = 4; ±SD). Significant differences from Col-0 were detected by Student’s *t*-test (\*\**p* < 0.01, \**p* < 0.05). C, Expression of *pEXPA7::GFP*, *pEXPA17::GFP*, and *pEXPA18::GFP* in lateral root primordia. Roots were stained with PI. Scale bars, 50 µm. D, RT-qPCR analysis of MYB93-regulated expansins in Col-0, *pXVE::CFP-gMYB93*/Col-0, *pXVE::CFP-gMYB93*/*myb93* #1, and *pXVE::CFP-gMYB93*/*myb93* #2 treated with 5 µM estradiol for 24 h. (n = 4; ±SD). Significant differences from Col-0 were detected by Student’s *t*-test (\*\**p* < 0.01, \**p* < 0.05).

### LTPG1 and LTPG2 were involved in RCC formation

We next investigated expression of *MYB93* and *EXPA17* in *ltpg1* and *ltpg2* mutants, and found comparable levels to those of Col-0 (Figure 7A). This indicated that the VLCFA signal regulating the MYB93 transcriptional network were functioned independently from LTPG. Considering the function of LTPGs in transporting VLCFAs out of the cell, and because VLCFAs are major RCC components, LTPG1 and LTPG2 were presumed to be involved in RCC formation, rather than in the VLCFA-mediated transcriptional network. Therefore, we examined RCC formation in *ltpg1* and *ltpg2* mutants via fluorol yellow (FY) staining, and observed that the number of stained LRPs in *ltpg1* and *ltpg2* plant was greatly reduced compared to that in Col-0. In addition, reduced stained was seen in *kcs1-5* mutants (Figure 7B). These results indicated VLCFA transport was important for LR development through RCC formation.

**Figure 7.**
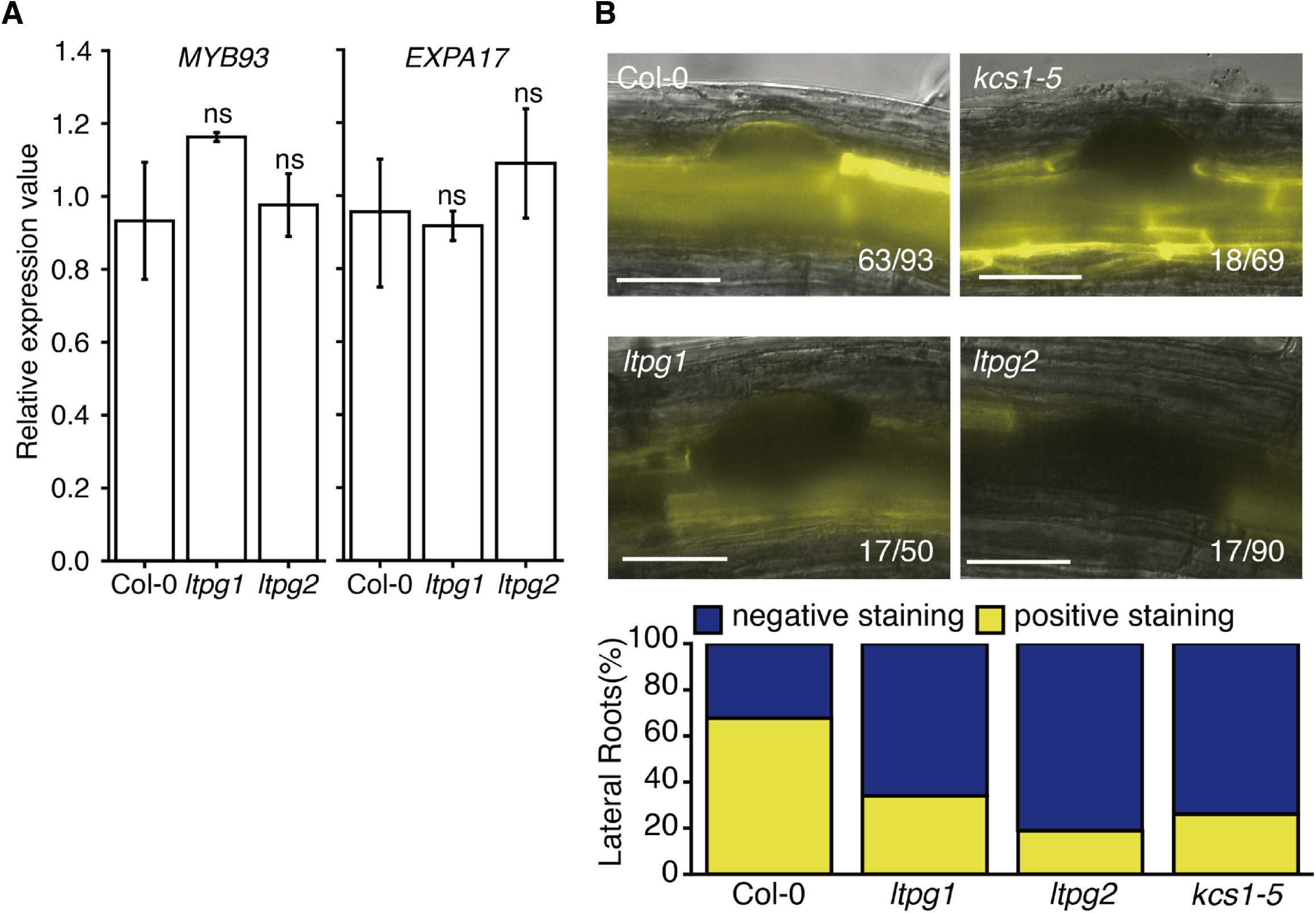
Root cuticle formation of LRPs of Col-0, *ltpg1* and *ltpg2*. A, RT-qPCR analysis of *MYB93* in seven-day-old whole roots of Col-0, *ltpg1*, and *ltpg2* plants (n = 4; ±SD). B, Fluorol yellow (FY088) staining of developing lateral roots in Col-0, *ltpg1*, *ltpg2*, and *kcs1-5* plants. Fractions in the figure indicate the number of individuals stained with FY088 (numerator: with staining; denominator, total number).

## Discussion

In this study, we elucidated part of the mechanism for LR development regulated by MYB93 in response to VLCFA content. Several recent studies have revealed that VLCFAs are important regulators for plant development. However, signaling behavior of VLCFAs remains poorly understood. Our findings offer to elucidate the role of VLCFAs in transcriptional network regulation. According to transcriptome analysis of *kcs1-5* mutants and expression analysis of exogenous C20, C22, and C24 treatment, the expression of *MYB93* was regulated by VLCFA levels. MYB93 is known as an early inhibitor of LR development, and *MYB93* is expressed in the LROC endodermis, where it is thought to regulate LRP development (Gibbs et al., 2014; Shukla et al., 2021). Exogenous auxin induces *MYB93* expression in the root basal meristem, where LRP development begins (Gibbs et al., 2014). Therefore, MYB93 is thought to function downstream of the first auxin signaling module to initiate LR production. In addition to this, DNN analysis revealed that MYB93 regulated late-stage LR development, especially LR emergence. The transition to late-stage LR development, from stage V to stage VI, was faster in *myb93* mutants than in wild type plants. This directly opposed the phenotype seen in *kcs1-5* mutants and Col-0 plants under cafenstrole treatment, which corresponded to *MYB93* overexpression in both Col-0 and *myb93* mutants. Because a lower level of VLCFA in *kcs1-5* was induced *MYB93*, the LR development phenotype indicated that *MYB93* was regulated by VLCFA and controlled late-stage LR development. Furthermore, in *kcs1-5*, the expression level of *MYB93* went down to the wild type level in three hours of exogenous VLCFA treatment. This strongly indicated that *MYB93* is a VLCFA responsive transcription factor that regulates LR development. RNAseq analysis of the LR development site in *myb93* mutants suggested that MYB93 repressed the expression of genes involved in cell wall organization. Since auxin-responsive genes were not significantly enriched in our data set, it is likely that auxin does not function downstream of MYB93. Among the MYB93-regulated genes, several expansin genes were detected. Expansins are known as the cell wall loosening enzymes (McQueen-Mason and Cosgrove, 1994; Cosgrove, 2005) and play variety roles for plant development such as hypocotyl elongation, root development, and pollen tube growth (Son et al., 2012; Ma et al., 2013; Lee and Kim, 2013; Lee et al., 2013; Liu et al., 2021). The detailed involvement of EXP17 in LR development has been reported for *Arabidopsis* (Lee and Kim, 2013, Lee et al., 2013). LR emergence is delayed in the *EXPA17* RNAi knockdown lines, and slightly increases under *EXPA17* oversxpression (Lee and Kim, 2013). This indicates that altering *EXPA17* expression can be used to control the emergence of LRs. In our study, *EXPA17* expression was upregulated in *myb93* mutants and downregulated in *MYB93* overexpressors. Accordingly, LR emergence was promoted in *myb93* mutants and suppressed in *MYB93* overexpressors. Moreover, *pEXPA17::GFP* was expressed in cortex LROCs, whereas *pMYB93::CFP-MYB93* expressed in endodermis LROCs. This could indicate that MYB93 repressed *EXPA17* expression in endodermis of LROCs. According to the cistrome database (http://neomorph.salk.edu/dap_web/pages/index.php), MYB93 can bind to the promoter region of *EXPA17*. This strongly indicates that MYB93 controls LR emergence by repressing *EXPA17* expression in endodermis of LROCs.

*LTPG1* and *LTPG2* were expressed specifically in the outermost layer of the LRP, and *ltpg1* and *ltpg2* mutants exhibited reduced LR numbers comparing to Col-0, especially emerged LRs. These results indicate that LTPG1 and LTPG2 are involved in LR development. LTPG is considered a VLCFA transporter from the endoplasmic reticulum to the surface of stems and leaves for wax cuticle synthesis (Debono et al., 2009; Kim et al., 2012). We previously reported that LTPG1 and LTPG2 may regulate root cell elongation via VLCFA transportation to the root tip elongation zone (Mabuchi et al., 2018). Moreover, it was recently found that defective RCC development leads delayed LR emergence (Berhin et al., 2019). This finding strongly indicates that VLCFAs, which are components of cuticle wax, play a critical role in LR development by forming RCCs. LTPG1 and LTPG2 may be involved in this process that transport VLCFA as substrates for RCC production. Accordingly, the expression patterns among *LTPG1*, *LTPG2*, *GPAT4*, and *GPAT8*, which are encoded for critical enzymes for cutin biosynthesis (Beisson et al., 2012), were quite similar in the LRP (Berhin et al., 2019). RCC-defective mutants, such as *gpat4*/*gpat8*, *dcr*, and *bdg* exhibit reduced LR emergence (Berhin et al., 2019). According to GC/MS analysis, VLCFA levels were comparable among wild type, *ltpg1*, and *ltpg2* mutants. This indicated that the reduction of LR numbers seen in *ltpg2* mutant may have been caused by defective transporting of VLCFA that resulted in lack of the RCC. Moreover, it has been reported that shoot cuticular layer structures in *ltpg1* and *ltpg2* mutants are altered and the total wax loads are reduced comparing Col-0 plants (Kim et al., 2012). This supported our hypothesis that LTPG1 and LTPG2 are involved in RCC formation. Additionally, FY staining of LRPs indicated reduced RCC in *ltpg1* and *ltpg2* mutants. In addition to *ltpg1* and *ltpg2*, *kcs1-5* mutants also showed reduced RCC formation in the LRP. This indicated that VLCFA regulated LRP development not only through transcriptional network signaling but also by constructing the RCC as a physiological barrier.

We began this study inferring that VLCFA was involved in LR development based on the expression patterns of *LTPG1* and *LTPG2* in the LRP. We were then able to identify part of the transcriptional network, which was regulated by MYB93, involved in LR development in response to root VLCFA levels. However, LTPG1 and LTPG2 were not involved in that transcriptional network, suggesting an additional regulatory system for LR development, likely involving RCC formation. Therefore, we have identified two important regulatory mechanisms of LRP development that are controlled by VLCFAs. Our work reveals the VLCFAs downstream gene regulatory network that control LR development. We have also found other putative VLCFA-responsive MYB transcription factors in addition to MYB93 in the *kcs1-5* RNAseq data set. In the near future, further functional analysis of these MYB transcription factors will reveal that VLCFAs are not just cellular components, but are signaling molecules in the transcriptional regulation of plant development.

## Materials and methods

### Plant materials and growth conditions

*Arabidopsis thaliana* Columbia-0 (Col-0) was used as the wild type. The T-DNA insertion line, *ltpg1* (CA878046), *ltpg2* (SALK_016947), *kcs1-5* (SALK_200839C), and *myb93* (SALK_131752) were obtained from the SALK collection and the seed stock center of the Arabidopsis Biological Resource Center. These mutants were genotyped using left-border primers on the T-DNA (LB), right-side primers on the genome (RP), and left-side primers on the genome (LP) (Supplemental Table S1). RP and LP primers were designed on the T-DNA express web site (http://signal.salk.edu/tdnaprimers.2.html).

All seeds were sterilized with 1% bleach and 0.05% Triton X-100 for 5 min, and then washed three times with sterilized water. Seeds were germinated on Murashige and Skoog (MS; FUJIFILM Wako Pure chemical, Osaka, Japan) medium supplemented with 1% sucrose and 1% agarose after two days at 4°C. Plants were grown vertically in a chamber (Panasonic, Osaka, Japan) at 22°C with a 16 h light/8 h dark cycle. For specific treatments, 10-day-old seedlings were transferred onto MS agarose plates containing 300 nM cafenstrole (FUJIFILM Wako pure chemical), 200 µM octadecanoic acid (C18; Tokyo Chemical Industry, Tokyo, Japan), 200 µM Arachidic acid (C20; Tokyo Chemical Industry), 200 µM behenic acid (C22; Tokyo Chemical Industry), or 200 µM lignoceric acid (C24; Tokyo Chemical Industry). For C20, C22, and C24 mixture treatment (VLCFA mix), seven-day-old seedlings were transferred onto MS agarose plates containing 200 µM each of arachidic acid, behenic acid, and lignoceric acid. For estradiol treatment, seven-day-old seedlings were transferred onto MS agarose plates containing 5 µM estradiol (FUJIFILM Wako pure chemical).

### Plasmid construction and plant transformation

Genomic DNA from Col-0 was used as the template for amplification of 2,981, 3,002, 1,619, and 3,003 bp upstream regions of *pMYB93*, *pAtEXPA7*, *pAtEXPA17*, and *pAtEXPA18,* respectively, for promoter cloning. One base of 5’ dA overhang was added to the PCR amplicon of *pMYB93* using Taq polymerase (Takara Bio Inc. Shiga, Japan), which was then cloned into pENTR5’-TOPO (Thermo Fischer Scientific, Waltham, MA, USA) and named pENTR-5’-*pMYB93*. To clone of *pAtEXPA7*, *pAtEXPA17*, and *pAtEXPA18,* the *pMYB93* region in pENTR-5’-*pMYB93* was replaced with *pAtEXPA7*, *pAtEXPA17*, and *pAtEXPA18* PCR amplicons, respectively, using NEBuilder^®^ HiFi DNA Assembly Cloning kit (New England Biolabs, Ipswich, MA, USA). For *LTPG1* and *LTPG2*, cDNA regions were amplified by PCR using the Col-0 root cDNA library as a template with a CACC sequence added just before each start codon for TOPO cloning. The CACC-cDNAs fragments were then cloned into pENTR/D-TOPO (Thermo Fisher Scientific). For *YFP-LTPG1* and *YFP-LTPG2* cloning, a *YFP* sequence was inserted just after putative endoplasmic reticulum signal peptide sequences in LTPG1 and LTPG2 using the homologous recombination Gibson Assembly system (New England Biolabs), according to Kim et al. (2012). YFP PCR amplicons were inserted into the pENTR/D-TOPO plasmids containing cDNA regions of *LTPG1* and *LTPG2,* 96 and 66 bp from the start codons of *LTPG1* and *LTPG2*, respectively. These PCR products were then subjected to homologous recombination using Gibson Assembly Master Mix, and YFP was inserted just after the signal sequences of *LTPG1* and *LTPG2*. The genomic region of *MYB93* was amplified by PCR using the Col-0 genome as a template, and the amplicon was inserted into the 3’ end of the pDONR201-CFP plasmid using the NEBuilder^®^ HiFi DNA Assembly Cloning kit.

For *pLTPG1::YFP-LTPG1* and *pLTPG2::YFP-LTPG2* constructs, the *LTPG1* and *LTPG2* (regions cloned by Mabuchi et al. [2018]) promoter regions containing pENTR5’-TOPO and *YFP-LTPG1*/ *YFP-LTPG2* containing pENTR/D-TOPO were cloned into R4pGWB501 (Nakagawa et al., 2008) using LR Clonase II (Thermo Fischer Scientific). For *pMYB93::CFP-MYB93*, pENTR-5’-*pMYB93* and *CFP-MYB93* containing pDONR201 were cloned into R4pGWB401 (Nakagawa et al., 2008) using LR Clonase II. For the *pXVE::CFP-MYB93* construct, *CFP-MYB93* containing pDONR201 was cloned into pMDC7 (Moore et al., 2006) using LR Clonase II. For *pEXPA7::GFP*, *pEXPA17::GFP*, and *pEXPA18::GFP*, the *EXPA7*, *EXPA17*, and *EXPA18* promoter regions, respectively, containing pENTR5’-TOPO were cloned into R4L1pGWB450 (Nakamura et al., 2009) using LR Clonase II. The resulting plasmids (*pLTPG1::YFP-LTPG1*, *pLTPG2::YFP-LTPG2*, *pMYB93::CFP-MYB93*, *pXVE::CFP-MYB93*, *pEXPA7::GFP*, *pEXPA17::GFP*, and *pEXPA18::GFP*) were transferred into *Agrobacterium tumefaciens* (C58C1 pMP90) cells and transformed into Col-0, *ltpg1*, *ltpg2*, and *myb93* mutants. The list of primers used in this study is provided in Supplemental Table S1.

### Quantitative real-time RT-PCR

RNA was isolated from whole roots of seven-day-old Col-0 treated with control MS medium, 200 µM VLCFA mix, and 300 µM cafenstrole for 1 d. *kcs1-5* mutants were treated with 200 µM C18, 200 µM C20, 200 µM C22, 200 µM C24, and 200 µM VLCFA mix for 1 d using the RNeasy Plant kit (QIAGEN, Hilden, Germany). For isolation of RNA from the bending site of roots, only the bending site of five-day-old roots that were gravistimulated for 46 h and microdissected (Tsukagoshi et al., 2010). The RNeasy micro kit (QIAGEN) was then used for RNA isolation. First-strand cDNA was synthesized using the ReverTra Ace qPCR RT Master Mix with gDNA Remover (TOYOBO Co., Ltd., Osaka, Japan). Quantitative real-time RT-PCR (RT-qPCR) was performed using the THUNDERBIRD SYBR qPCR Mix (TOYOBO) on a real-time PCR Eco system (PCRmax, Stone Staffordshire, UK). The primers used in this study are listed in Supplemental Table S1. RT-qPCR efficiency and CT value were determined using the standard curves for each primer set. Efficiency-corrected transcript values of three biological replicates for all samples were used to determine relative expression values. Each value was normalized against the level of PDF2 (Czechowski, et al, 2005).

### RNAseq experiments

cDNA libraries were generated from 1 µg of whole-root RNA from 13-day-old Col-0 and *kcs1-5* plants, and 100 ng of bending site RNA samples from five-day-old Col-0 and *myb93* plants using the NEBnext Ultra II RNA Library Prep kit (New England Biolabs) following manufacturer protocols. Both ends of the cDNA libraries were sequenced for 60 cycles using a paired-end module with an Illumina NextSeq 500 platform (Illumina, San Diego, CA, USA). Two biological replicates were conducted for each experiment.

### RNAseq data analysis

The short read sequencing results were mapped to the Arabidopsis genome (TAIR10: www.arabidopsis.org/) using Bowtie software (Langmead et al., 2009). These datasets were normalized, then False Discovery Rate (FDR) and Fold Change (FC) were calculated using the edgeR package for R (Robinson et al., 2010). We used an FDR of q < 0.05 as the cut-off to determine differentially expressed genes between Col-0 and *ksc1-5* and between Col-0 and *myb93*. The data were deposited in the DNA Data Bank of Japan (DDBJ) sequence Read Archive (DRA) (https://www.ddbj.nig.ac.jp/index-e.html) with the accession number DRA013878. Enriched GO categories were analyzed using the agriGO v2.0 website (Tian et al., 2017; http://systemsbiology.cau.edu.cn/agriGOv2/).

### Phenotypic and microscopic analysis

Lateral root numbers in whole roots of 13-day-old seedlings were counted using an Olympus SZX7 microscope (Olympus, Tokyo, Japan). Stage classification was determined according to Malamy and Benfey (1997). For treatment with cafenstrole (300 nM), 10-day-old seedlings grown in MS medium were transferred to media containing cafenstrole, and the seedlings were grown 3 d before counting LRs.

For LR induction by gravistimulation, vertically-grown, five-day-old Col-0, *kcs1-5*, and *myb93* seedlings were rotated 90 degrees. For cafenstrole (300 nM), cafenstrole (300 nM) + VLCFA mix (200 µM each), and VLCFA mix (200 µM each) treatment, four-day-old Col-0 and *kcs1-5* seedlings were transferred to media containing relevant compounds for 24 h, then rotated 90 degrees. As for *pXVE::CFP-MYB93*, before rotating 90 degrees, four-day-old *pXVE::CFP-MYB93* seedlings were pretreated with 5 µM estradiol for 24 h. LR stages were analyzed after 20, 32, and 46 h of gravistimulation using a Leica DMI 6000B-AFC (Leica Camera, Wetzlar, Germany). Roots were cleared using a clearing solution (a mixture of 20 g of chloral hydrate in 1 mL of glycerol and 6 mL of water). Experiments were repeated independently three times (n = 20 in each experiment).

For RCC visualization, Fluorol Yellow 088 staining was performed according to the method described by Berhin et al (2019). Eight-day-old Col-0, *kcs1-5*, *ltpg1*, and *ltpg2* seedlings were stained with Fluorol Yellow 088 (0.01% in methanol; Santa Cruz Biotechnology, Dallas, TX, USA) for 3 days at 4°C. Root specimens were then counterstained with aniline blue (0.5% in water) for minimum of 1 h at 25°C, and imaged with a Leica DMI 6000B-AFC microscope (Leica).

Laser scanning confocal microscopy for *pLTPG1::YFP-LTPG1*, *pLTPG2::YFP-LTPG2*, *pMYB93::CFP-gMYB93*, *pEXPA7::GFP*, *pEXPA17::GFP*, and *pEXPA18::GFP* was performed with a Leica SP8 system (Leica) on propidium iodide (PI; FUJIFILM Wako pure chemical) stained roots. Roots were stained with PI in a 10 µg mL^-1^ dilution in water for 3-5 min, with 448 nm excitation and 460-510 nm emission for CFP, 488 nm excitation and 490-543 nm emission for YFP, and 555 nm excitation with 580-680 nm emission for PI. Image assembly was performed using LAS X software (Leica).

For time-lapse imaging, a Lab-Tek Chambered Coverglass w/cvr (Thermo Fisher Scientific) was used as described previously (Mabuchi et al., 2018). Four-day-old seedlings in MS medium were rotated 90 degrees for six hours to induce LR formation by gravistimulation, then the seedlings were placed on the chambered cover glass. For treatments with cafenstrole (300 nM), VLCFA mix (200 µM each of C20, C22, and C24), and estradiol (5 µM), three-day-old seedlings were pretreated with these compounds for 24 h and then gravistimulated for 6 h, and placed on the chambered cover glass in media containing treatments. The plants were imaged with DMI 6000B-AFC microscope and images were captured by the LAS X system every 30 min for 66 h.

### Training data and machine learning for DNN

To produce training data, images of LRP stages Ⅰ-Ⅶ taken with the Leica DMI 6000B-AFC system were manually classified. The number of training images for each stage was 636 for stage 0, 283 for stage I, 247 for stage II, 356 for stage III, 366 for stage IV, 813 for stage V, 305 for stage VI, and 1,107 for stage VII. The LRP classification model was based on the ResNet50 pretrained by ImageNet, with the last layer modified. We used the Keras Library for implementation. Inputs and outputs shape was 500 x 500 x 3 and 1 respectively (He et al. 2016). All networks were trained with the Adam optimizer to minimize the mean squared error (Supplemental Figure S2A). The model at the 87 epoch of the training phase was used for prediction of test images (Supplemental Figure S2B) because the MAE values of the training and validation data were similar (Supplemental Figure S2C). The MAE value of the validation data showed the minimum possible value, and the generalization performance was considered high.

For the training and validation image dataset, we prepared total of 4,123 time-lapse images and individual LRP images and classified them into LRP staged 0 to VII. DNN training and validation were performed on 3,923 and 200 images, respectively, which were randomly assigned in the Python script. The test image dataset was created by classifying a single human into LRP stages 0 to VII using 358 time-lapse images acquired independently of those used for DNNs training and validation. The 358 test images were classified four times each by three different unbiassed individuals (for a total of 12 LRP classification results), and the MAE values for manual classification were calculated by comparing the differences in the stages obtained from all of 66 combinations of the 12 LRP classification results. The MAE value from the DNN classification was also calculated using the same 358 test LRP images. The accuracy of DNN model was evaluated by comparing the resulting MAE value with that of the manual classifications (Supplemental Figure S2, C, D, and E). Generated time series data of LRP stages (Supplemental Figure S2F) were approximated through a quadratic function. The source code generated during this study can be found at https://github.com/yuta-Bio/xception_LRP_classification.

### Fatty acid extraction and analysis

The extraction of fatty acids was performed according to described in Patel et al. (2018) with some modifications. Briefly, Col-0 roots grown in MS medium for 10 days and transferred to e control MS medium or MS medium with 300 nM cafenstrole 3 d. *ksc1-5*, *ltpg1*, and *ltpg2* roots were grown in MS medium for 13 d. Root samples (500 mg) of each were corrected, and frozen, and ground. The ground powder was transferred to chloroform-methanol (1:2) solution, then washed with an equal volume of phosphate buffered saline (pH 7.5) to obtain lipid samples. To prepare fatty acid methyl esters (FAMEs), lipid samples were transmethylated with NaOH (v/v, 1% in methanol), then heated at 55°C for 15 min. Before adding NaOH, 0.025 mg of nonadecanoic acid (Tokyo Chemical Industry) was added as the internal standard for subsequent gas chromatography-mass spectrometry (GC-MS) analysis. Then, 2 mL of methanolic HCL (v/v, 5%) was added and the tubes were heated at 55 °C for an additional 15 min. FAMEs were extracted in aqueous hexane (v/v, 1:2), lyophilized, and dissolved in 120 µL hexane. GC-MS was performed at 70 eV using a GCMS-QP2010 (Shimadzu Corporation, Kyoto, Japan) instrument equipped with a 30 m fused silica column (DB-5; J&W Scientific, Inc., Folsom, CA, USA). The oven temperature was programmed to ramp from 50°C to 300°C at 3.5°C min^-1^, with an injection temperature of 280°C. FAMEs were identified by comparing their respective GC retention times and mass fragmentation patterns with those of authentic standards. FAMEs in each sample were normalized using peak areas of nonadecanoic acid, and were then quantified using corresponding standards (C4-C24 even carbon saturated FAMEs [Sigma-Aldrich]).

### Statistical analysis

All statistical analysis were conducted using Microsoft Excel or R programs. Details of analysis are given in the figure legends.

## Accession numbers

Sequence data from this article can be found in the Arabidopsis Initiative under the following accession numbers: *LTPG1* (At1g27950), *LTPG2* (At3g43720), *MYB93* (At1g34670), *EXPA7* (At1g12560), *EXPA17* (At4g01630), and *EXPA18* (At1g62980).

## Supplemental data

**Supplemental Figure S1.** Fatty acids composition of Col-0, *ltpg1*, and *ltpg2* roots.

**Supplemental Figure S2.** Validation of DNN analysis.

**Supplemental Figure S3.** Expression level of *MYB93* in Col-0, *myb93*, *pXVE::CFP-gMYB93*/Col-0, and *pXVE::CFP-gMYB93*/*myb93*.

**Supplemental Figure S4.** GFP expression of *pEXPA7::GFP*, *pEXPA17::GFP*, and *pEXPA18::GFP* in whole roots.

**Supplemental Table S1.** Primers used in this study.

**Supplemental Data Set S1.** Significantly differentially expressed genes between Col-0 and *kcs1-5*.

**Supplemental Data Set S2.** Significantly differentially expressed genes between Col-0 and *myb93*.

## Acknowledgments

We thank Dr. W. Busch for comments on the manuscript, and the Arabidopsis Biological Resource Center for providing seeds.

## Author Contributions

H.T. conceived the project. H.T. and A.M. designed the project. Y.U. and K.H. performed DNN analysis. Y.U., S.K., S.S., and H.T. performed plant genotyping, plasmid construction, plant transformation, and RNA preparation. Y.U., S.S., and H.T. performed RT-qPCR. Y.U. and S.K. performed RNA-seq library preparation. T.S. performed RNA sequencing. Y.U., S.T., and H.T. analyzed RNAseq data. Y.U., S.K., T.O., and Y.K. performed LR stage analysis. Y.U. and K.M. performed confocal microscopy. Y.U. performed time-lapse imaging. S.K., H.K., and M.S. performed GC-MS experiment and analyzed the data. Y.U. and H.T. wrote the paper and produced the figures.

## Conflict of interest statement

None decleared.

**Supplemental Figure S1.** Fatty acids composition of Col-0, *ltpg1*, and *ltpg2* roots. Data are represented as mean ± SD of three biological replicates. Fatty acids composition of Col-0 was determined from data in Figure 2B. Significant differences from Col-0 of each stage were detected by Student’s *t*-test (\*\**p* < 0.01, \**p* < 0.05).

**Supplemental Figure S2.** Validation of DNN analysis. Comparison of manual and machine classification for each lateral root stage. Three individuals and a machine each performed lateral root primordium stage classification on 358 images, and the accuracy was indicated as mean absolute error (MAE). A, Schematic view of DNN model. B, Transition of training (light blue line) and validation (orange line) loss during DNN learning. Y and X axis indicated loss (mean squared error), and epochs, respectively. (C) Comparison of MAE between DNN and human lateral root primordium classification accuracy. *p*-value was 0.3887, using Welch’s test. D, MAE of training, validation, and test DNN classification. E, Representative DNN classification of lateral root primordium stage images. The numbers in each image indicate the lateral root primordium stage classified by DNN. F, Averaged raw data of lateral root primordium stage transition of Col-0 (orange line; n = 29), *kcs1-5* (blue line; n = 24), *kcs1-5* treated with VLCFA mix (red line; n = 12), and Col-0 treated with cafenstrole (green line; n = 19). Shaded area indicates the 95 % confidence interval.

**Supplemental Figure S3.** Expression level of *MYB93* in Col-0, *myb93*, *pXVE::CFP-gMYB93*/Col-0, and *pXVE::CFP-gMYB93*/*myb93*. RT-qPCR analysis of *MYB93* in seven-day-old whole roots of Col-0, *myb93*, and *pXVE::CFP-gMYB93*/Col-0, *pXVE::CFP-gMYB93*/*myb93* #1, and *pXVE::CFP-gMYB93*/*myb93* #2 plants treated with 5 µM estradiol for 0 and 24 h (n = 4; ±SD). Significant differences from Col-0 were detected by Student’s *t*-test (\*\**p* < 0.01).

**Supplemental Figure S4.** GFP expression of *pEXPA7::GFP*, *pEXPA17::GFP*, and *pEXPA18::GFP* in whole roots. Seven-day-old whole roots of *pEXPA7::GFP*, *pEXPA17::GFP*, and *pEXPA18::GFP*. Roots were stained with propidium iodide (PI). A, C, and E; merged images of GFP and PI, B, D, and F; GFP fluorescent images. Scale bars, 150 µm.

**Supplemental Table S1.** Primers used in this study.

## References

1. Bach L, Michaelson LV, Haslam R, Bellec Y, Gissot L, Marion J, Da Costa M, Boutin JP, Miquel M, Tellier F, et al. (2008) The very-long-chain hydroxy fatty acyl-CoA dehydratase PASTICCINO2 is essential and limiting for plant development. Proc Natl Acad Sci USA 105: 14727–14731

2. Bach L, Faure JD (2010) Role of very-long-chain fatty acids in plant development, when chain length does matter. C R Biol 333: 361–370

3. Bach L, Gissot L, Marion J, Tellier F, Moreau P, Satiat-Jeunemaître B, Palauqui JC, Napier JA, Faure JD (2011) Very-long-chain fatty acids are required for cell plate formation during cytokinesis in Arabidopsis thaliana. J Cell Sci 124: 3223–3234

4. Batsale M, Bahammou D, Fouillen L, Mongrand S, Joubès J, Domergue F (2021) Biosynthesis and Functions of Very-Long-Chain Fatty Acids in the Responses of Plants to Abiotic and Biotic Stresses. Cells 10: 1284

5. Beaudoin F, Wu X, Li F, Haslam RP, Markham JE, Zheng H, Napier JA, Kunst L (2009) Functional characterization of the Arabidopsis beta-ketoacyl-coenzyme A reductase candidates of the fatty acid elongase. Plant Physiol 150: 1174–1191

6. Beisson F, Li-Beisson Y, Pollard M (2012) Solving the puzzles of cutin and suberin polymer biosynthesis. Curr Opin Plant Biol 15: 329–337

7. Berhin A, de Bellis D, Franke RB, Buono RA, Nowack MK, Nawrath C (2019) The Root Cap Cuticle: A Cell Wall Structure for Seedling Establishment and Lateral Root Formation. Cell 176: 1367–1378.e8

8. Cosgrove DJ (2005) Growth of the plant cell wall. Nat Rev Mol Cell Biol 6: 850–861.

9. Czechowski T, Stitt M, Altmann T, Udvardi MK, Scheible WR (2005) Genome-wide identification and testing of superior reference genes for transcript normalization in Arabidopsis. Plant Physiol 139: 5–17

10. De Bigault Du Granrut A, Cacas JL (2016) How Very-Long-Chain Fatty Acids Could Signal Stressful Conditions in Plants? Front Plant Sci 7: 1490

11. Debono A, Yeats TH, Rose JK, Bird D, Jetter R, Kunst L, Samuels L (2009) Arabidopsis LTPG is a glycosylphosphatidylinositol-anchored lipid transfer protein required for export of lipids to the plant surface. Plant Cell 21: 1230–1238

12. Devaiah SP, Roth MR, Baughman E, Li M, Tamura P, Jeannotte R, Welti R, Wang X (2006) Quantitative profiling of polar glycerolipid species from organs of wild-type Arabidopsis and a phospholipase Dalpha1 knockout mutant. Phytochemistry 67: 1907–1924

13. Du Y, Scheres B (2018) Lateral root formation and the multiple roles of auxin. J Exp Bot 69: 155–167

14. Dubrovsky JG, Sauer M, Napsucialy-Mendivil S, Ivanchenko MG, Friml J, Shishkova S, Celenza J, Benková E (2008) Auxin acts as a local morphogenetic trigger to specify lateral root founder cells. Proc Natl Acad Sci USA 105: 8790–8794

15. Edstam MM, Edqvist J (2014) Involvement of GPI-anchored lipid transfer proteins in the development of seed coats and pollen in Arabidopsis thaliana. Physiol Plant 152: 32–42

16. Gibbs DJ, Voß U, Harding SA, Fannon J, Moody LA, Yamada E, Swarup K, Nibau C, Bassel GW, Choudhary A, et al. (2014) AtMYB93 is a novel negative regulator of lateral root development in Arabidopsis. New Phytol 203: 1194–1207

17. He K, Zhang X, Ren S, Sun J (2016) Deep Residual Learning for Image Recognition. 2016 IEEE Conference on Computer Vision and Pattern Recognition (CVPR) pp.770–778. doi: 10.1109/CVPR.2016.90.

18. Joubès J, Raffaele S, Bourdenx B, Garcia C, Laroche-Traineau J, Moreau P, Domergue F, Lessire R (2008) The VLCFA elongase gene family in Arabidopsis thaliana: phylogenetic analysis, 3D modelling and expression profiling. Plant Mol Biol 67: 547–566

19. Kim H, Lee SB, Kim HJ, Min MK, Hwang I, Suh MC (2012) Characterization of glycosylphosphatidylinositol-anchored lipid transfer protein 2 (LTPG2) and overlapping function between LTPG/LTPG1 and LTPG2 in cuticular wax export or accumulation in Arabidopsis thaliana. Plant Cell Physiol 53: 1391–1403

20. Kumpf RP, Shi CL, Larrieu A, Stø IM, Butenko MA, Péret B, Riiser SR, Bennett MJ, Aalen RB (2013) Floral organ abscission peptide IDA and its HAE/HSL2 receptors control cell separation during lateral root emergence. Proc Natl Acad Sci USA 110: 5235–5240

21. Langmead B, Trapnell C, Pop M, Salzberg SL (2009) Ultrafast and memory-efficient alignment of short DNA sequences to the human genome. Genome Biol 10: R25

22. Lee SB, Go YS, Bae HJ, Park JH, Cho SH, Cho HJ, Lee DS, Park OK, Hwang I, Suh MC (2009) Disruption of glycosylphosphatidylinositol-anchored lipid transfer protein gene altered cuticular lipid composition, increased plastoglobules, and enhanced susceptibility to infection by the fungal pathogen Alternaria brassicicola. Plant Physiol 150: 42–54

23. Lee HW, Kim J (2013) EXPANSINA17 up-regulated by LBD18/ASL20 promotes lateral root formation during the auxin response. Plant Cell Physiol 54: 1600–1611

24. Lee HW, Kim MJ, Kim NY, Lee SH, Kim J (2013) LBD18 acts as a transcriptional activator that directly binds to the EXPANSIN14 promoter in promoting lateral root emergence of Arabidopsis. Plant J 73: 212–224

25. Liu W, Xu L, Lin H, Cao J (2021) Two Expansin Genes, AtEXPA4 and AtEXPB5, Are Redundantly Required for Pollen Tube Growth and AtEXPA4 Is Involved in Primary Root Elongation in Arabidopsis thaliana. Genes (Basel) 12: 249

26. Lv B, Wei K, Hu K, Tian T, Zhang F, Yu Z, Zhang D, Su Y, Sang Y, Zhang X, et al. (2021) MPK14-mediated auxin signaling controls lateral root development via ERF13-regulated very-long-chain fatty acid biosynthesis. Mol Plant 14: 285–297

27. Ma N, Wang Y, Qiu S, Kang Z, Che S, Wang G, Huang J (2013) Overexpression of OsEXPA8, a root-specific gene, improves rice growth and root system architecture by facilitating cell extension. PLoS One 8: e75997

28. Mabuchi K, Maki H, Itaya T, Suzuki T, Nomoto M, Sakaoka S, Morikami A, Higashiyama T, Tada Y, Busch W, et al. (2018) MYB30 links ROS signaling, root cell elongation, and plant immune responses. Proc Natl Acad Sci USA 115: E4710–E4719

29. Malamy JE, Benfey PN (1997) Organization and cell differentiation in lateral roots of Arabidopsis thaliana. Development 124: 33–44

30. McQueen-Mason S, Cosgrove DJ (1994) Disruption of hydrogen bonding between plant cell wall polymers by proteins that induce wall extension. Proc Natl Acad Sci USA 91: 6574–6578

31. Markham JE, Lynch DV, Napier JA, Dunn TM, Cahoon EB (2013) Plant sphingolipids: function follows form. Curr Opin Plant Biol 16: 350–357

32. Millar AA, Kunst L (1997) Very-long-chain fatty acid biosynthesis is controlled through the expression and specificity of the condensing enzyme. Plant J 12: 121–131

33. Molino D, Van der Giessen E, Gissot L, Hématy K, Marion J, Barthelemy J, Bellec Y, Vernhettes S, Satiat-Jeunemaître B, Galli T, et al. (2014) Inhibition of very long acyl chain sphingolipid synthesis modifies membrane dynamics during plant cytokinesis. Biochim Biophys Acta 1842: 1422–1430

34. Moore I, Samalova M, Kurup S (2006) Transactivated and chemically inducible gene expression in plants. Plant J 45: 651–683

35. Morineau C, Gissot L, Bellec Y, Hematy K, Tellier F, Renne C, Haslam R, Beaudoin F, Napier J, Faure JD (2016) Dual Fatty Acid Elongase Complex Interactions in Arabidopsis. PLoS One 11: e0160631

36. Motte H, Beeckman T (2019) The evolution of root branching: increasing the level of plasticity. J Exp Bot 70: 785–793

37. Nakagawa T, Nakamura S, Tanaka K, Kawamukai M, Suzuki T, Nakamura K, Kimura T, Ishiguro S (2008) Development of R4 gateway binary vectors (R4pGWB) enabling high-throughput promoter swapping for plant research. Biosci Biotechnol Biochem 72: 624–629

38. Nakamura S, Nakao A, Kawamukai M, Kimura T, Ishiguro S, Nakagawa T (2009) Development of Gateway binary vectors, R4L1pGWBs, for promoter analysis in higher plants. Biosci Biotechnol Biochem 73: 2556–2559

39. Nobusawa T, Okushima Y, Nagata N, Kojima M, Sakakibara H, Umeda M (2013) Synthesis of very-long-chain fatty acids in the epidermis controls plant organ growth by restricting cell proliferation. PLoS Biol 11: e1001531

40. Parizot B, Laplaze L, Ricaud L, Boucheron-Dubuisson E, Bayle V, Bonke M, De Smet I, Poethig SR, Helariutta Y, Haseloff J, et al. (2007) Diarch symmetry of the vascular bundle in Arabidopsis root encompasses the pericycle and is reflected in distich lateral root initiation. Plant Physiol 146: 140–148

41. Patel MK, Das S, Thakur JK (2018) GC-MS-Based Analysis of Methanol: Chloroform-extracted Fatty Acids from Plant Tissues. Bio Protoc 8: e3014

42. Robinson MD, McCarthy DJ, Smyth GK (2010) edgeR: A bioconductor package for differential expression analysis of digital gene expression data. Bioinformatics 26: 139–140

43. Roudier F, Gissot L, Beaudoin F, Haslam R, Michaelson L, Marion J, Molino D, Lima A, Bach L, Morin H, et al. (2010) Very-long-chain fatty acids are involved in polar auxin transport and developmental patterning in Arabidopsis. Plant Cell 22: 364–375

44. Shang B, Xu C, Zhang X, Cao H, Xin W, Hu Y (2016) Very-long-chain fatty acids restrict regeneration capacity by confining pericycle competence for callus formation in Arabidopsis. Proc Natl Acad Sci USA 113: 5101–5106

45. Shukla V, Han JP, Cleard F, Barberon M (2021) Suberin plasticity to developmental and exogenous cues is regulated by a set of MYB transcription factors. Proc Natl Acad Sci USA 118: e2101730118

46. Son SH, Chang SC, Park CH, Kim SK (2012) Ethylene negatively regulates EXPA5 expression in Arabidopsis thaliana. Physiol Plant 144: 254–262

47. Swarup K, Benková E, Swarup R, Casimiro I, Péret B, Yang Y, Parry G, Nielsen E, De Smet I, Vanneste S, et al. (2008) The auxin influx carrier LAX3 promotes lateral root emergence. Nat Cell Biol 10: 946–954

48. Teixeria JAS, ten Tusscher KH (2019) The Systems Biology of Lateral Root Formation: Connecting the Dots. Mol Plant 12: 784–803

49. Tian T, Liu Y, Yan H, You Q, Yi X, Du Z, Xu W, Su Z (2017) agriGO v2.0: a GO analysis toolkit for the agricultural community, 2017 update. Nucleic Acids Res 45: W122–W129

50. Todd J, Post-Beittenmiller D, Jaworski JG (1999) KCS1 encodes a fatty acid elongase 3-ketoacyl-CoA synthase affecting wax biosynthesis in Arabidopsis thaliana. Plant J 17: 119–130

51. Trenkamp S, Martin W, Tietjen K (2004) Specific and differential inhibition of very long-chain fatty acid elongases from Arabidopsis thaliana by different herbicides. Proc Natl Acad Sci USA 101: 11903–11908

52. Trinh DC, Lavenus J, Goh T, Boutté Y, Drogue Q, Vaissayre V, Tellier F, Lucas M, Voß U, Gantet P, et al. (2019) PUCHI regulates very long chain fatty acid biosynthesis during lateral root and callus formation. Proc Natl Acad Sci USA 116: 14325–14330

53. Tsukagoshi H, Busch W, Benfey PN (2010) Transcriptional regulation of ROS controls transition from proliferation to differentiation in the root. Cell 143: 606–616

54. Vermeer JEM, von Wangenheim D, Barberon M, Lee Y, Stelzer EHK, Maizel A, Geldner N (2014) A spatial accommodation by neighboring cells is required for organ initiation in Arabidopsis. Science 343: 178–183

55. Zheng H, Rowland O, Kunst L (2005) Disruptions of the Arabidopsis Enoyl-CoA reductase gene reveal an essential role for very-long-chain fatty acid synthesis in cell expansion during plant morphogenesis. Plant Cell 17: 1467–1481

